# Differentiable simulation to develop molecular dynamics force fields for disordered proteins

**DOI:** 10.1101/2023.08.29.555352

**Authors:** Joe G Greener

## Abstract

Implicit solvent force fields are computationally efficient but can be unsuitable for running molecular dynamics on disordered proteins. Here I improve the a99SB-*disp* force field and the GBNeck2 implicit solvent model to better describe disordered proteins. Differentiable molecular simulations with 5 ns trajectories are used to jointly optimise 108 parameters to better match explicit solvent trajectories. Simulations with the improved force field better reproduce the radius of gyration and secondary structure content seen in experiments, whilst showing slightly degraded performance on folded proteins and protein complexes. The force field, called GB99dms, reproduces the results of a small molecule binding study and improves agreement to experiment for the aggregation of amyloid peptides. GB99dms, which can be used in OpenMM, is available at https://github.com/greener-group/GB99dms. This work is the first to show that gradients can be obtained directly from nanosecond-length differentiable simulations of biomolecules and highlights the effectiveness of this approach to training whole force fields to match desired properties.

## Introduction

Molecular dynamics (MD) simulations have helped us to understand how molecules move [1, 2], and will only become more important as computers get faster and innovative machine learning approaches are developed [3, 4]. There are two main issues with MD: the force fields used to describe atomic interactions lack accuracy, and sampling beyond the microsecond scale is computationally prohibitive. When simulating a biomolecular system it is usually the solute that is of interest, so implicit solvent models can be used to replace solvent molecules with a continuous medium [5, 6]. This speeds up simulations by reducing the number of atoms and by giving faster exploration of conformational space due to the lack of friction with solvent, with speedups of up to 100x over explicit solvent [7]. Despite the lack of interactions between the solute and individual solvent molecules being a fundamental limitation of implicit solvent models, they are regularly used across a range of biomolecular simulations [8] and have been the target of machine learning approaches [9, 10]. Small proteins can be folded in GPU-days with implicit solvent models and enhanced sampling [11].

Major inaccuracies of implicit solvent models include the tendency to overcompact disordered proteins into rigid, often α-helical structures, the tendency to cause any pair of proteins to bind strongly, and the poor secondary structure match to experiments for peptides [12, 13, 14, 15]. There has been considerable recent effort to alleviate similar, and less severe, problems for explicit solvent force fields [16, 17, 18], resulting in a better match to experimental data for both folded and disordered proteins [19, 20, 21, 22]. This forms part of a wider attempt to use data to improve force fields [23, 24, 25, 26, 27]. However there has been less attention on improving implicit solvent models in this area [28], with notable exceptions being the ABSINTH model [29] and coarse-grained approaches that combine multiple atoms into one site [30, 31]. This is a missed opportunity: despite their fundamental limitations, implicit solvent models need not perform as poorly as they do on disordered systems. In fact they are especially well-suited to studying these systems, as they are not slowed by the large solvent boxes required for explicit solvent simulations and can be used to probe slow events such as protein aggregation [32, 33] with the increased conformational sampling resulting from low viscosity. Whereas explicit solvent force fields have been improved in tandem with associated water models [34, 35, 36, 37], this is rarely the case for implicit solvent models [12, 38], suggesting that adjusting both at the same time could lead to improvements. Whilst it is possible and even desirable to train force fields using only quantum mechanical data [39, 40, 41, 42, 43], matching to structural properties as well can be a route to higher accuracy, particularly for biomolecules where a wealth of structural data is available.

In this study I modify the parameters of an existing force field and implicit solvent model to better describe disordered proteins and protein aggregation, whilst retaining acceptable performance on folded proteins. The technique of differentiable molecular simulation (DMS) is used to modify many force field parameters at the same time. This emerging technique allows structural and dynamic properties to be targeted to parameterise whole force fields using automatic differentiation (AD) [44]. This is part of the paradigm of differentiable programming [45] in which AD, more commonly associated with training neural networks, is used to obtain gradients from arbitrary algorithms. DMS has previously been used to train a coarse-grained force field for proteins from scratch [46], to learn pairwise potentials [47, 48], for enhanced sampling [49], to predict protein structure [50] and to explore statistical physics models [51]. Dedicated software packages such as Jax MD [52], TorchMD [53] and DMFF [54] have been developed specifically for differentiable simulations. The Molly.jl software developed as part of this work provides a flexible and fast option for DMS and for MD more broadly. The combined force field and implicit solvent model presented here, called GB99dms, is available to the community and is easy to run from OpenMM. Here it is shown for the first time that DMS can be used to train force fields over nanosecond-length simulations.

## Results

There are a number of recent software packages designed for DMS [52, 53, 54]. However all were either lacking features for simulating proteins or did not have the performance required to do AD on simulations of millions of steps. Hence the capabilities of the Julia [55, 56] package Molly.jl were expanded to carry out training simulations for this work. This package is a pure Julia implementation of MD compatible with biomolecules and DMS which implements various integrators and allows easy definition of custom interactions. Code is run on the GPU using kernels written in Julia with CUDA.jl [57, 58]. Gradients are computed using Zygote.jl [59] and Enzyme.jl [60, 61]. Together these allow gradients to be computed through arbitrary code on the GPU, including complicated algorithms such as the force calculation of GBNeck2. Reverse-mode AD is used as it has constant compute time with respect to the number of parameters. However, broadcasted functions do use an efficient combined forward and reverse approach [62].

I choose to start from the a99SB-*disp* force field [21] since it has been developed for both folded and disordered proteins, and the GBNeck2 implicit solvent model [63] since it shows good performance on folded proteins [11]. These have been developed over many years using both quantum mechanical and experimental data, and here we seek to improve the combination of the models for disordered proteins. GBNeck2 is a Generalized Born approach that approximates the exact Poisson-Boltzmann equation describing the electrostatic environment of a solute in a solvent. Generalized Born methods model the solute as a set of spheres with a different dielectric constant to the external solvent. The “neck” correction improves the prediction of the molecular surface, which is used when determining the Born radii [64]. The a99SB-*disp* modified backbone O-H interaction term is not used, as it was later found to not impact the ability to fit quantum mechanical data [22]. Since some atom types in the force field appear rarely in the training data, parameters are modified for 16 common atom types: CA, CT, C, C8, C9, N, N3, O, O2, OH, H, H1, HA, HC, HO and HP. This gives 108 parameters to change: partial charge scaling, Lennard-Jones (LJ) σ and LJ ε for the 16 atom types (46 non-zero parameters); torsion *k* values for 13 common proper torsions (33); LJ and Coulomb 1-4 interaction scalings (2); GBNeck2 atom radii (6); GBNeck2 atom parameters (12); GBNeck2 screening parameters (4); and the GBNeck2 parameters neck cutoff, neck scale, offset, probe radius and surface area factor (5). To maintain the overall charge of residues the partial charges are not changed directly, instead a charge scaling is learned - see the methods. Torsion phases, which in a99SB-*disp* are all symmetrical around 0 degrees apart from backbone φ and ψ, are not modified as this has questionable physical validity.

8 small proteins ranging in size from 19 to 39 residues, 4 folded and 4 disordered, are used for training. These are shown in Figure 1B. The folded proteins are Trp-cage (PDB ID 2JOF), BBA (1FME), GTT (2F21) and NTL9 (2HBA) and contain both α-helices and β-sheets. The disordered proteins are Htt-1-19, a slightly helical intrinsically disordered protein (IDP) derived from huntingtin’s N-terminus; histatin-5; and the N-terminal and C-terminal halves of ACTR, split up to avoid excessive computational demands during training. At each epoch of training, each protein is simulated using a Langevin integrator with the current force field parameters for 5 ns (5 million steps). Residue-residue distances are periodically recorded and used at the end of the simulation to calculate the mean and standard deviation for each residue-residue distance over the simulation. Since explicit solvent simulations with the a99SB-*disp* force field and its corresponding water model seem to accurately describe the properties of both folded and disordered proteins [21], I train to match the residue-residue distances in 2 µs simulations of the training proteins with this force field. Previous approaches to developing implicit solvent force fields have also matched to explicit solvent data [65].

**Figure 1.**
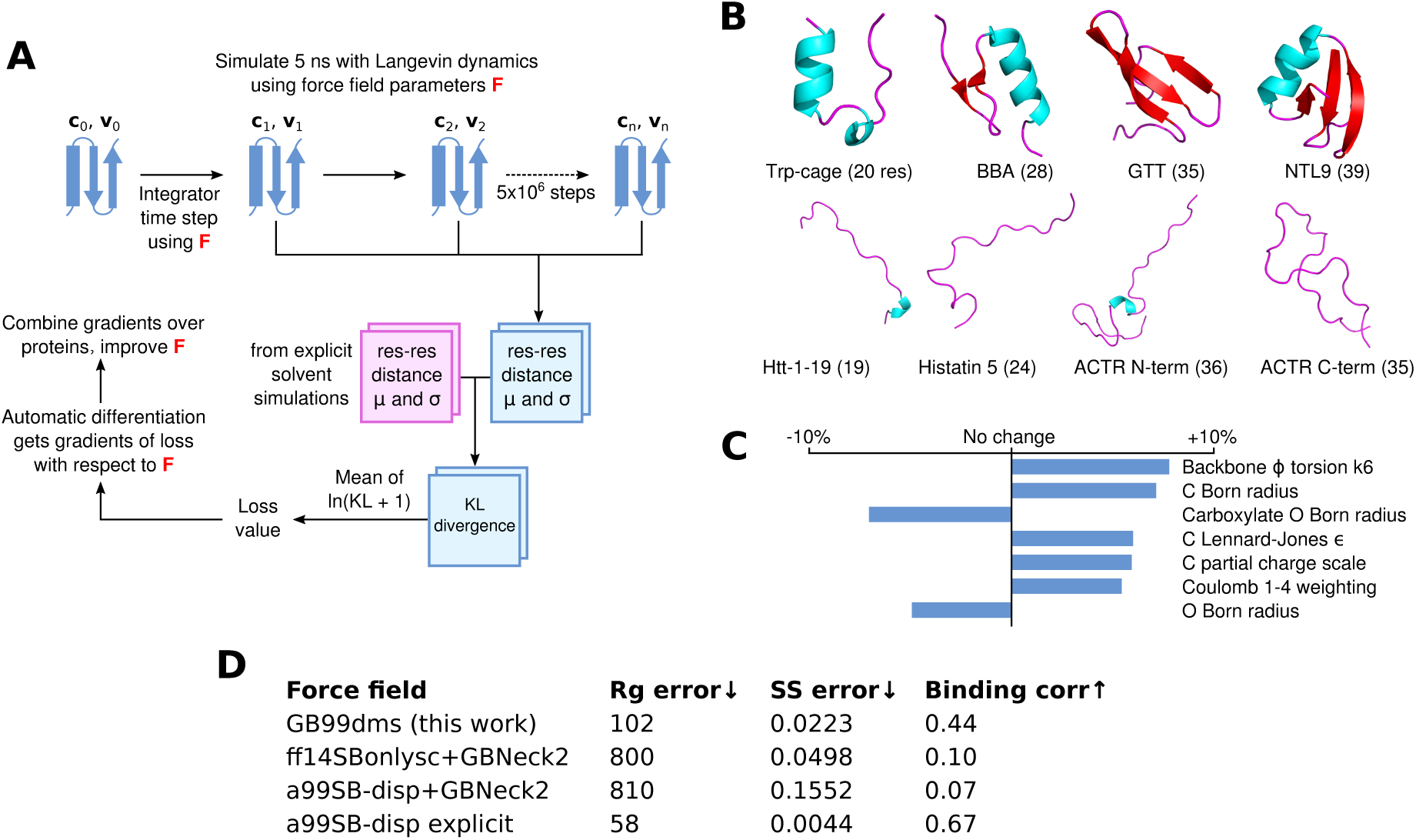
Differentiable molecular simulation to improve an implicit solvent force field. (A) For each protein a simulation is run with Langevin dynamics and the mean (µ) and standard deviation (σ) of the Cα residue-residue distances are calculated. These are compared using the KL divergence to corresponding values from reference explicit solvent simulations to obtain a loss value. AD is then used to obtain gradients of the loss with respect to the force field parameters, which can be used to improve the force field. (B) The 8 proteins used for training, consisting of 4 folded proteins (top) and 4 disordered proteins (bottom). (C) The 7 parameters that change by at least 4.5% in absolute value from the starting values in a99SB-*disp*+GBNeck2. (D) Summary of numerical results from the paper. Rg error is the sum of squared total residuals between the experimental and simulation radius of gyration for each protein as shown in Figure 2A, lower is better. SS error is the α-helical fraction error as shown in Figure 3B, lower is better. Binding corr is the correlation between simulation contact probabilities and NMR chemical shift perturbation data for α-synuclein and fasudil as described in the results, higher is better.

The Kullback-Leibler (KL) divergence is calculated in both directions between the training and reference simulations, and averaged over residues. This gives a loss value; since the simulation is implemented in AD-compatible software, the gradients of the loss with respect to the force field parameters can be calculated. These are combined across the training proteins and used to change the force field parameters, before the next epoch is started. Simulations of the folded proteins start from the PDB coordinates and a single repeat is run. For the disordered proteins two repeats are run, starting from different snapshots of the reference explicit solvent simulations. Each 5 ns simulation takes 15-24 hours on a GeForce RTX 2080 Ti GPU depending on the size of the protein. The parameters after 5 epochs of training are used, meaning training takes 5 days on 12 GPUs (the repeats for disordered proteins give 12 simulations per epoch). This overall training process is shown in Figure 1A.

The parameters of the trained force field, named GB99dms, are similar to the starting parameters, with only 19 out of 108 parameters changing by more than 3% in absolute value. Staying close to the starting values was a deliberate attempt to find a point in the high-dimensional parameter space that works across a variety of protein systems but remains similar to the well-studied parameters available currently [66]. The direction of parameter changes are generally consistent across epochs of training, as shown in Figure S1. The parameters that change the most are shown in Figure 1C and all parameters are listed in Table S1. The Born radius parameters for carbonyl O and backbone amide H decrease, indicating less screening from solvent, though the Born radius parameters for C and H increase. The LJ σ parameters for N and C increase, indicating a larger interaction distance for these atoms. The LJ ε parameters increase for N and C but decrease for HC and CT, indicating a shift in strength of LJ interactions between atom types. Small changes are made to torsion parameters; as expected due to the required changes in secondary structure preferences, backbone φ and ψ change the most. Partial charges do not change much, except C which becomes more positive (0.597 to 0.625 for backbone carbonyl C in alanine). The Coulomb and LJ 1-4 interaction weightings for atoms separated by 3 bonds both increase, indicating less shielding from non-bonded forces for nearby atoms. Whilst the bonded/non-bonded parameters and implicit solvent parameters of this force field could be used separately in future, this was not tested and it is recommended that they are used together as they were trained.

GB99dms was compared to existing force fields on a set of proteins not homologous to those used for training. I compare to the combination of Amber ff14SBonlysc [11, 68] and GBNeck2 [63] used successfully to fold proteins [11], the combination of a99SB-*disp* [21] and GBNeck2 used at the start of training in this work, and available explicit solvent a99SB-*disp* data [21]. First, performance is assessed on 8 IDPs ranging in size from 40 to 140 residues. This is the set used in Robustelli et al. 2018 [21], excluding ACTR which is used here for training. For each protein 3 simulations of 2 µs were run for each force field, with an initial burn-in period of 0.5 µs starting from an extended conformation being discarded. All validation simulations were run with a Langevin collision frequency γ of 1 ps^-1^ to increase conformational sampling [11]. As shown in Figure 2 and Figure 1D the radius of gyration (*R_g_*) using GB99dms matches experimental data [21] and observed scaling laws [67] considerably better than the existing implicit solvent force fields across a range of protein sizes. Explicit solvent a99SB-*disp* still matches better, indicating that the accuracy of the reference data used during training is not the factor limiting improvement. Figure 3 and Figure 2C indicate that this is partly due to existing force fields giving structures that are too α-helical and inflexible, and Figure 2C shows that the experimental *R_g_* may be sampled with GB99dms even when the mean simulation *R_g_* is different. The overall α-helical content of IDPs also matches experiment better with GB99dms than with existing implicit solvent force fields, as shown in Figure 3B-C. The error is 0.022 for GB99dms compared to 0.050 for ff14SBonlysc and 0.155 for a99SB-*disp*. There is still some discrepancy between GB99dms and experiment, for example in the location of α-helices in Ntail and PaaA2, in line with the variety of secondary structure propensities shown previously by explicit solvent force fields for these IDPs [21].

**Figure 2.**
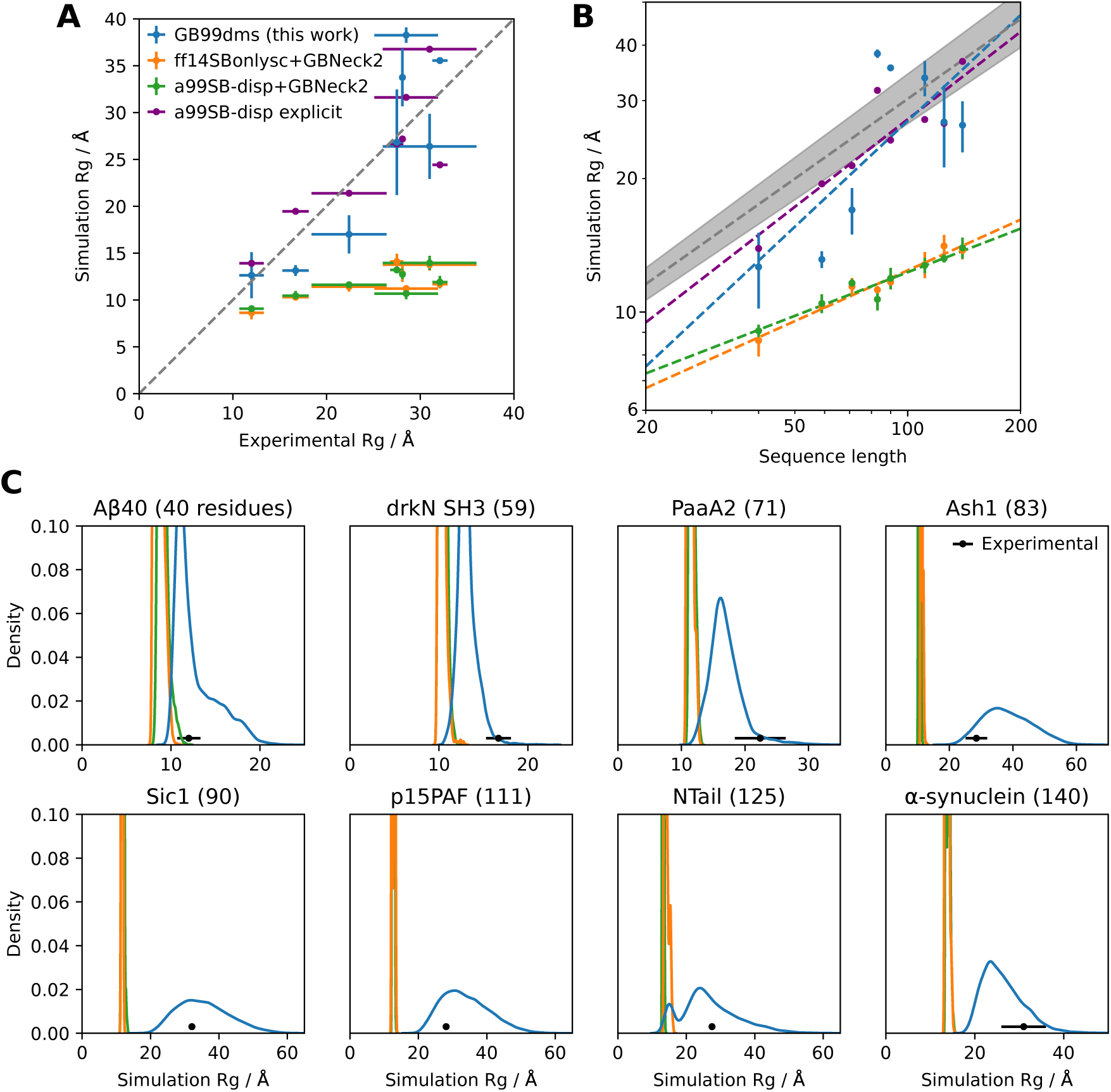
Radius of gyration of IDPs with different force fields. Experimental values for *R_g_*and simulation *R_g_* values for explicit solvent a99SB-*disp* with its corresponding water model are taken from Robustelli et al. 2018 [21]. (A) Comparison of simulation and experimental *R_g_*. Each point (excepting a99SB-*disp* explicit) is the mean *R_g_* over 3 simulations of 2 µs, with an initial burn-in period of 0.5 µs starting from an extended conformation being discarded for each simulation. The error bars in the x direction represent uncertainty in the experimental value and error bars in the y direction represent 95% confidence intervals of the mean calculated from the standard error of the mean across the 3 simulations. The dotted line represents equal experimental and simulation values. Summary data is shown in Figure 1D. (B) Comparison of simulation *R_g_* to sequence length for the same data. The dotted grey line represents the experimentally observed power law relationship *R_G_* = *R*_0_*N^v^* with *v* = 0.598 *±* 0.028 and *R*_0_ = 1.927 ^°^A [67]. The shaded grey area represents the 95% confidence interval of *v*. This power law relationship is for chemically denatured proteins and was mainly fit on proteins longer than 50 residues. The best fit *v* values from the simulation data for GB99dms, ff14SBonlysc+GBNeck2, a99SB-*disp*+GBNeck2 and a99SB-*disp* explicit are 0.793, 0.380, 0.327 and 0.656 respectively. (C) Distributions of simulation *R_g_* for the same data.

**Figure 3.**
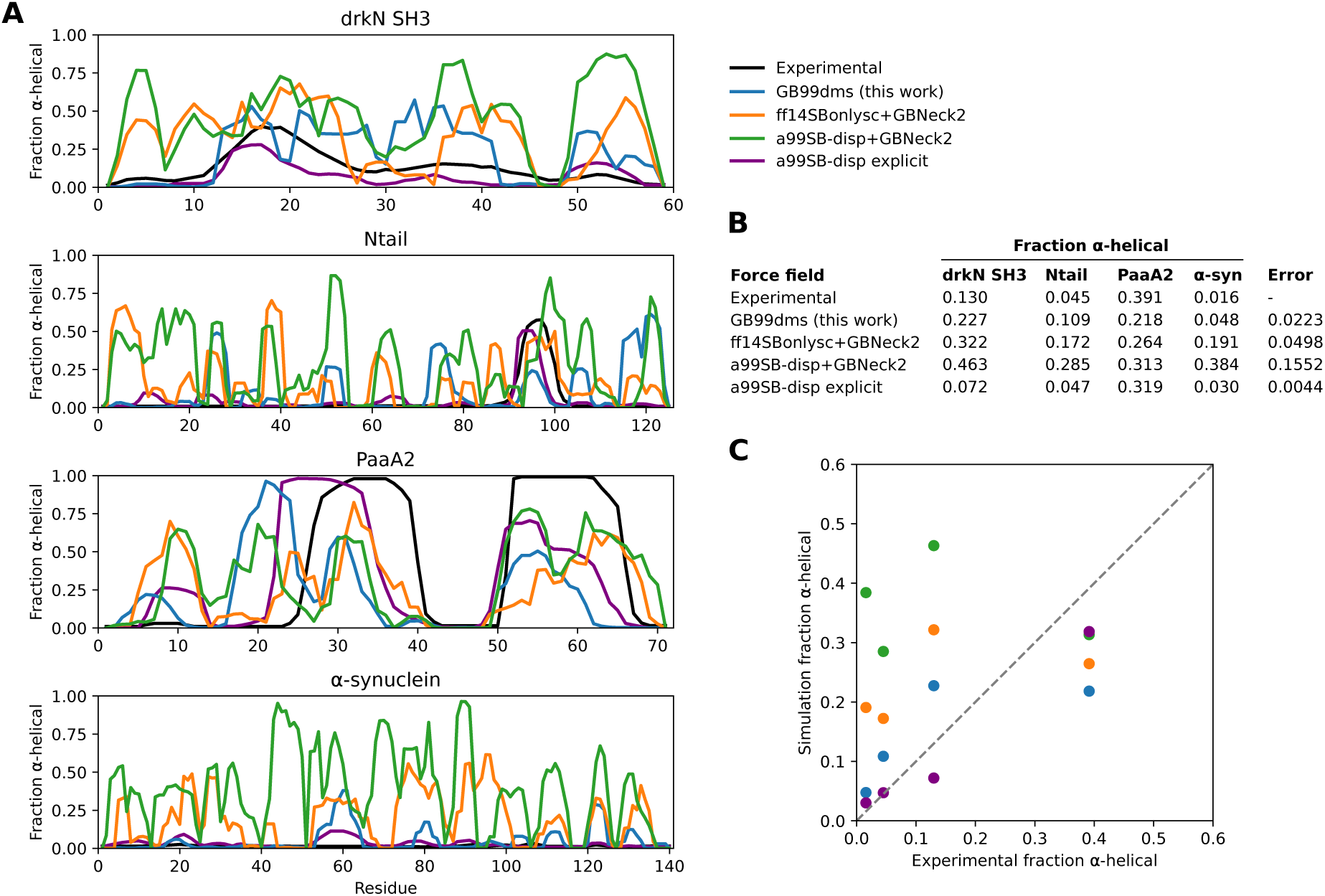
Secondary structure content of IDPs with different force fields. Experimental values for α-helical fraction and simulation values for explicit solvent a99SB-*disp* with its corresponding water model are taken from Robustelli et al. 2018 [21]. (A) The α-helical fraction for each residue of 4 IDPs with different force fields. The simulations are the same as in Figure 2. (B) A table showing the mean α-helical fraction across residues for each protein and force field. An overall error value is also calculated per force field as the sum of squared total residuals between the experimental and simulation mean α-helical fractions for each protein. (C) A plot of the simulation mean α-helical fractions against experimental values. The dotted line represents equal experimental and simulation values.

Having established improved performance on IDPs, it is important to test whether performance has degraded for folded proteins. 3 simulations of 2 µs for each force field were run on the 4 folded proteins in Robustelli et al. 2018 [21], ranging in size from 56 to 129 residues: GB3 (PDB ID 1P7E), ubiquitin (1D3Z), hen egg white lysozyme (HEWL, 6LYZ) and bovine pancreatic trypsin inhibitor (BPTI, 5PTI). Figure 4A shows the root-mean-square deviation (RMSD) to the native structure across the trajectory for each simulation. GB3 and ubiquitin show similar stability over the simulations with all 3 force fields. HEWL and BPTI show less stability with GB99dms. For HEWL 3 of the α-helices and the β-sheet lose their secondary structure, though the overall tertiary structure remains the same. For BPTI the β-sheet remains intact but the α-helices lose their secondary structure and the loops show significant flexibility. The difficulty of balancing secondary structure preferences in implicit solvent force fields has been established previously [13].

**Figure 4.**
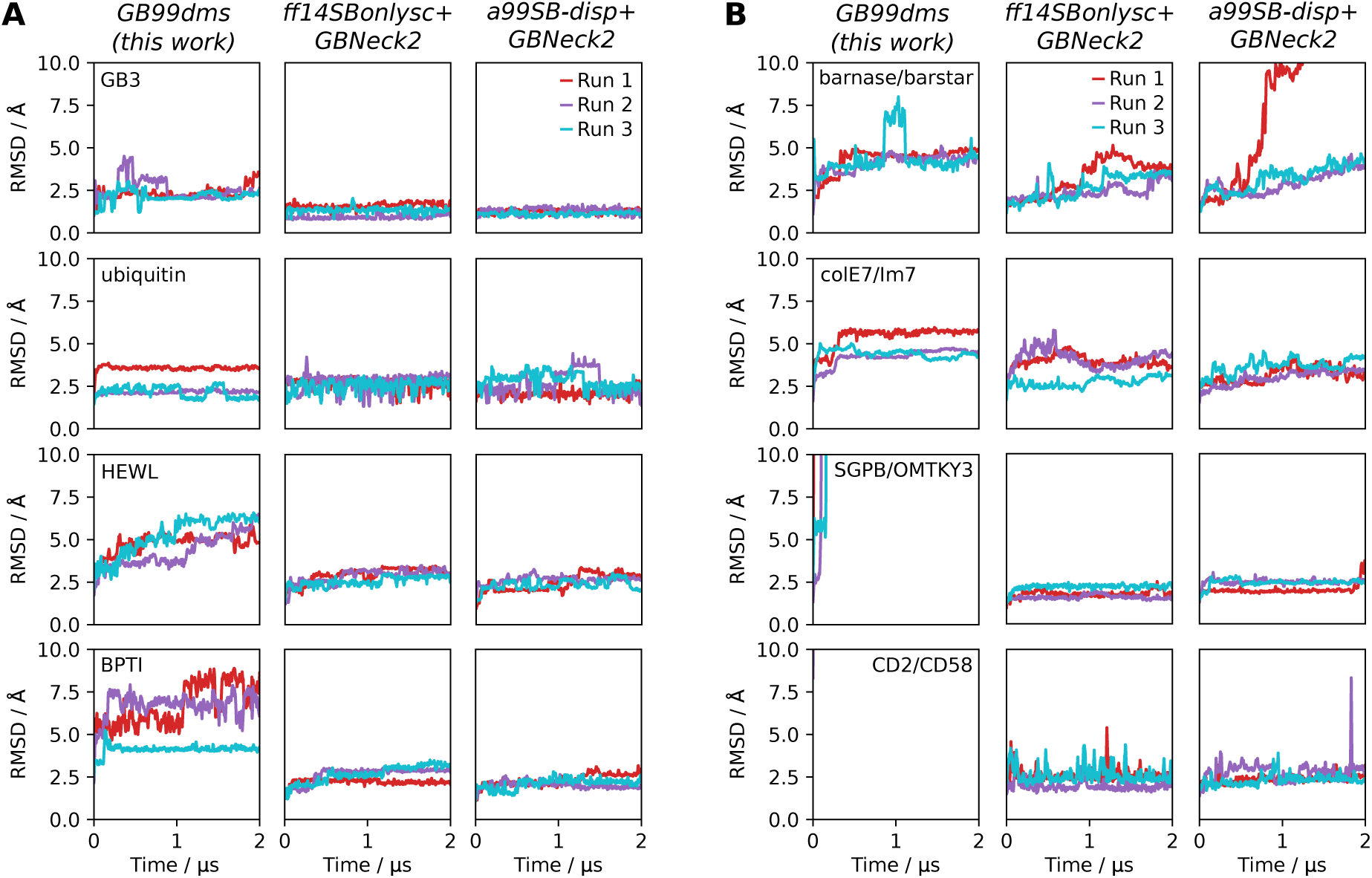
The behaviour of folded proteins and protein complexes with implicit solvent force fields. RMSD to the PDB starting structure is shown over 3 x 2 µs simulations. Snapshots are recorded every 0.5 ns and the mean RMSD of a window extending 10 snapshots either side of the given snapshot is shown. (A) Behaviour of 4 folded proteins. (B) Behaviour of 4 protein dimers. Dissociation occurs within 1 ns for CD2/CD58. Dissociation is not expected for any of the dimers on this time scale.

**Figure 5.**
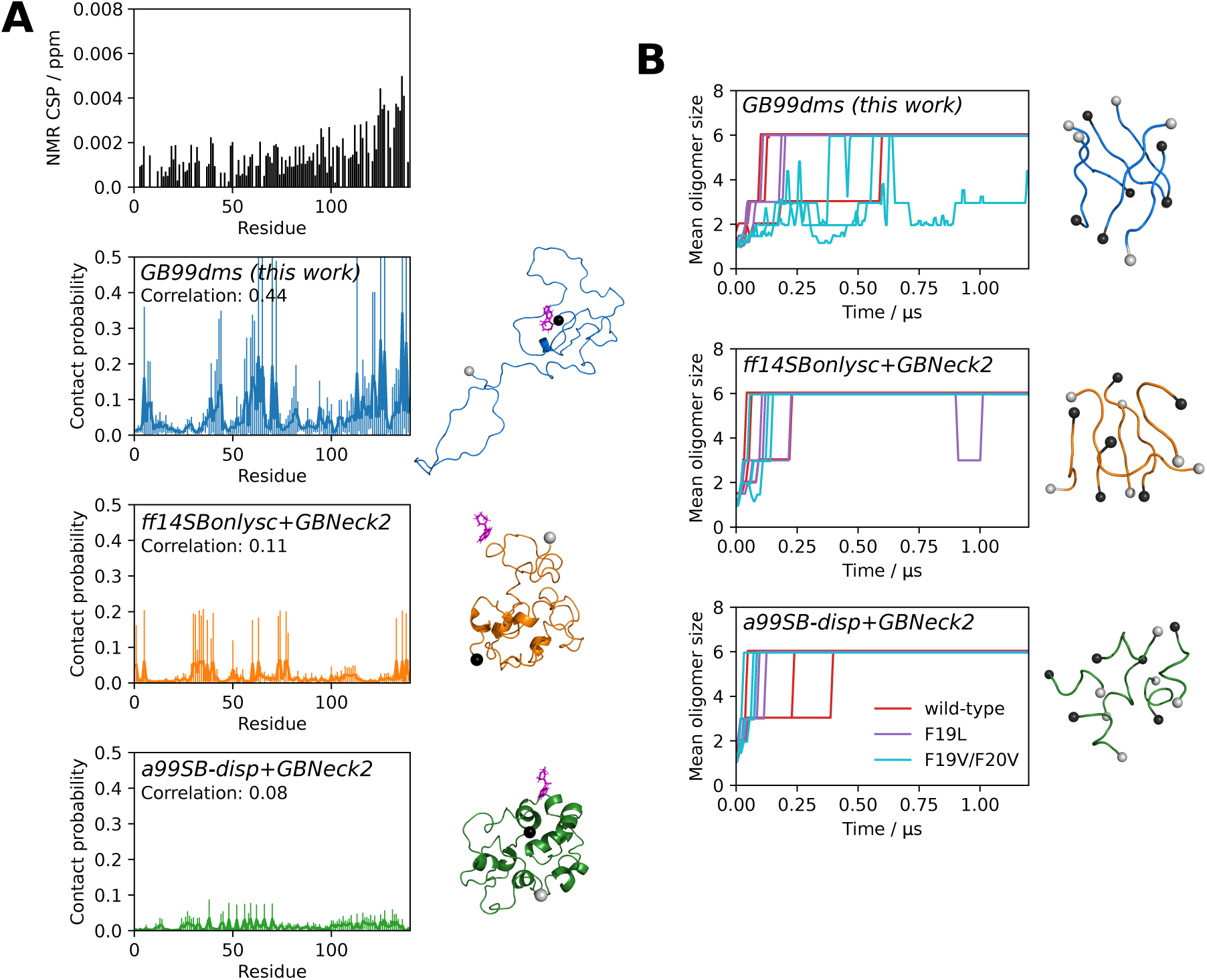
Ligand binding and amyloid aggregation with implicit solvent force fields. Grey and black spheres on structures represent the N- and C-termini respectively. (A) The interaction of α-synuclein and the small molecule fasudil. The NMR CSP data from Robustelli et al. 2022 [70] is shown. The fraction of the time each residue is in contact with fasudil over 5 x 2 µs simulations is shown for each force field, along with a snapshot where fasudil is in contact with α-synuclein. The error bars for each residue represent 95% confidence intervals of the mean calculated from the standard error of the mean across the 5 simulations. The Pearson correlation coefficient between the contact probabilities and the NMR CSP data is also shown. (B) Oligomerisation of 6 x Aβ_16-22_. Three capped peptides are studied: KLVFFAE (wild-type), F19L (faster aggregation) and F19V/F20V (no aggregation). The oligomer size over 3 x 2 µs simulations is shown for each sequence and force field. Repeats are shown separately, with the plots truncated at 1.2 µs as there is little change beyond this time. Snapshots are recorded every 0.5 ns and the mean oligomer size of a window extending 10 snapshots either side of the given snapshot is shown. The final oligomer from the first wild-type repeat is also shown.

Performance is also assessed on 4 medium-sized protein dimers from Piana et al. 2020 [22] ranging in size from 197 to 235 total residues: barnase/barstar (PDB ID 1X1X), colE7/Im7 (7CEI), SGPB/OMTKY3 (3SGB) and CD2/CD58 (1QA9). The dimers were simulated with no periodic boundaries since the intention was to explore initial unbinding events, which should not occur on the time scales used here. There has been less work on developing and assessing implicit solvent force fields for protein complexes, but here it is found that the 4 dimers are generally stable under the two existing force fields. Figure 4B indicates that with GB99dms barnase/barstar and colE7/Im7 seem stable, though SGPB/OMTKY3 and CD2/CD58 dissociate quickly. The dissociating complexes have less favourable experimental association free energy than those that remain bound [22]. With a collision frequency γ of 91 ps^-1^ more representative of the viscous drag of water [69], SGPB/OMTKY3 remains bound at 5 ^°^A after 2 µs RMSD but CD2/CD58 still dissociates. The difficulty of improving performance on IDPs whilst not allowing protein complexes to dissociate has also been encountered with explicit solvent force fields [21, 22]. Overall GB99dms has slightly degraded performance on folded proteins, though could still be useful for studying systems containing a mix of folded and disordered proteins if the stability and structural properties of the folded proteins with GB99dms are verified initially.

MD simulations can be used to study the interactions of small molecules with IDPs, assisting in the difficult process of finding drugs to target the many IDPs implicated in disease. For example, a recent study [70] used a99SB-*disp* to carry out 1.5 ms of explicit solvent simulation of α-synuclein with the small molecule fasudil. α-synuclein is associated with Parkinson’s disease and fasudil has been shown to delay α-synuclein aggregation [71]. The contact probability over the simulation of each residue with fasudil was shown to correlate with NMR chemical shift perturbation (CSP) data, allowing an interpretation to be made of how the drug affects the protein. CSPs are sensitive to changes in the local environment of each backbone amide bond and a higher CSP indicates protein-ligand interaction near that residue. I carried out similar simulations, using the faster sampling of implicit solvent to compare 5 simulations of 2 µs with the NMR data and significantly longer explicit solvent simulations from Robustelli et al. 2022 [70]. Figure 5A shows that GB99dms reproduces a similar profile to the NMR data. There is elevated contact probability in the residue 121-140 C-terminal region, peaks around residues 5-9/39-44/59-72/121-127/136-138, and Y136 has the highest contact probability. In contrast, ff14SBonlysc shows blocks of interacting residues consistent with fasudil interacting with the surface of a compact, inflexible structure and a99SB-*disp* shows consistent and lower interaction throughout the protein. The Pearson correlation coefficients between the contact probabilities and the NMR chemical shift perturbation data are 0.44, 0.11 and 0.08 for GB99dms, ff14SBonlysc and a99SB-*disp* respectively, compared to 0.67 for the explicit solvent simulations [70]. This indicates that GB99dms could be useful for assessing small molecule binding to IDPs during drug discovery. Though it is unusual to use periodic boundary conditions with implicit solvent - one advantage of implicit solvent is not having to deal with boundaries - it is possible since the solvent behaviour is unrelated to the boundary.

Finally, I investigate the behaviour of aggregating peptides with GB99dms. It has previously been shown that the oligomerisation behaviour under simulation of the 7-residue aggregating core of amyloid beta (Aβ), Aβ_16-22_, depends on the force field and does not always reproduce the effect of mutations [72, 73]. Here I study the oligomerisation of 6 capped Aβ_16-22_ peptides in a periodic box. I simulate 3 x 2 µs with each force field for 3 peptide sequences: the wild-type KLVFFAE, the F19L mutant which aggregates faster than wild-type, and the F19V/F20V double mutant which does not aggregate [73]. All force fields show oligomerisation of all 6 peptides within 2 µs. As shown in Figure 5B, GB99dms reproduces best the observed behaviour of the sequences: on average F19L forms oligomers fastest and F19V/F20V slowest. The wild-type forms oligomers faster than F19L for ff14SBonlysc and F19V/F20V forms oligomers at a similar speed to wild-type for ff14SBonlysc and a99SB-*disp*. The final oligomers adopt extended monomer conformations for GB99dms reminiscent of Aβ fibrils, whereas for ff14SBonlysc and particularly a99SB-*disp* the conformations are more α-helical. These results show that GB99dms could be used to study amyloid aggregation at scale. One advantage of implicit solvent is that the periodic box size, and hence the effective concentration of Aβ, can be changed without adding more atoms. Currently, high concentration is a limitation of many MD studies of aggregation [32].

## Discussion

Recently much effort has gone into developing machine learning interatomic potentials (MLIPs), which show quantum mechanical-level accuracy at faster speeds. Whilst these are promising, I believe that the techniques of machine learning such as AD can also be used to improve molecular mechanics force fields by effective optimisation of their large parameter spaces [74, 66]. This retains the advantages of interpretability, robustness and speed given by molecular mechanics force fields over MLIPs. The smooth nature of these force fields - at least compared to the hard potentials used in domains such as robotics simulations - makes them well-suited to differentiable simulation. An advantage of DMS over force-matching approaches is that targeting global structural properties alleviates issues when simulations reach states not seen in their training data, a problem that can lead to a lack of stability for MLIPs as errors accumulate during simulation [75]. Another advantage is that existing experimental data can potentially be used [46, 47], reducing the large amount of quantum mechanical data required to train accurate and transferable MLIPs. Most biomolecular force fields in wide use today have been developed using a combination of quantum mechanical and condensed phase experimental data, and DMS provides a route to fit all force field parameters to experimental data. In the long run it may be possible to forgo experimental data and train fast and accurate molecular mechanics force fields solely on quantum mechanical data [39, 40, 42], but until then approaches that improve force fields using various observed properties will be valuable. The popular ForceBalance method [23] is also able to target both quantum mechanical and experimental data, and DMS could enhance this approach by replacing the costly finite difference step used to obtaining gradients. Differentiable trajectory reweighting [76] has explored this direction as well.

The simulations of 5 million steps carried out in this work are the longest differentiable molecular simulations to date and indicate that even longer simulations are possible. The next step is to train an all-atom explicit solvent force field using DMS to match experimental data such as NMR constraints alongside existing training approaches to match quantum mechanical data. This will require improvements in computational speed, as well as differentiable implementations of algorithms such as bond constraints and Ewald summation that could run into the limitations of AD [77]. The flexibility of the approach allows it to be combined with other recent advances such as continuous atom typing with graph neural networks [78] and exploration of different functional forms for non-bonded interactions [79]. One promising approach, Time Machine (https://github.com/proteneer/timemachine), aims to use DMS for drug discovery.

A question surrounding DMS is whether accurate gradients can be obtained through long MD simulations. Gradients could explode or vanish, and there are also concerns about error propagation over long, chaotic simulations [80, 49, 50, 81]. Here, I find that the Langevin integrator is effective at propagating gradients using reverse-mode AD. By contrast, simulations in the NVE ensemble were not found to produce stable gradients. The friction and stochastic noise applied to every atom at every step in Langevin dynamics likely provides a regularisation effect that helps prevent gradient explosion but does not lead to vanishing gradients [82]. This stochasticity, along with the random starting velocities, means that the gradients are a sample over a distribution. When repeating runs, around 80% of the paired parameter gradients have the same sign and the Pearson correlation coefficient of paired parameter gradients is over 0.85. This is shown in Table S2. Good correlation is also found when comparing simulations run with a 1 fs and a 0.5 fs time step and when adding noise to the starting parameters. Adjoint sensitivity methods provide another way to obtain gradients through simulations [83, 84, 48], but are often unstable and have had less development than reverse-mode AD. Here I find that the “simple” approach of using AD on the integrator works well. It may be possible to find speedups due to the iterative and reversible nature of molecular simulation [83].

Another question is whether the computational overhead of calculating the gradients via AD makes it worthwhile compared to using a black box approach such as finite differencing. The Julia code used here is currently 100-1000x slower on the GPU when gradients are required than heavily optimised non-differentiable codes such as OpenMM [85] and Gromacs [86]. However the gradients for all parameters are calculated to numerical accuracy in one go, whereas a number of runs would be required per gradient when using finite differencing. It seems that the current case of optimising 108 parameters is around the crossover point, and optimising any fewer parameters would have been easier with finite differencing. As DMS code becomes faster and more parameters are included for training, for example more atom types or torsion CMAP potentials, the advantage of DMS will become clearer. One direction of future work is to improve the performance of the Julia code significantly using more advanced GPU kernels [57] and further use of the Enzyme AD framework [60, 61]. It remains an open question how close in performance a differentiable implementation can get to a non-differentiable one. Molly.jl will be useful for exploring these questions, and for training the next generation of force fields that are transferable, reproducible and fast.

## Methods

The a99SB-*disp* force field was manually converted from Desmond/Gromacs files to the OpenMM XML force field format with the modified backbone O-H interaction term omitted. Training simulations were carried out with Molly.jl, which is available along with documentation at https://github.com/JuliaMolSim/Molly.jl and will be described fully in a future publication. Training simulations were carried out for 5 ns using a Langevin integrator with a γ of 0.1 ps^-1^, a 1 fs time step, a temperature of 300 K and no distance cutoff for non-bonded interactions. A Debye-Hückel screening parameter κ of 0.7 nm^-1^ was used [87]. This is roughly equivalent at 300 K to a salt concentration of 100 mM, which is similar to that used when simulating biomolecules under physiological conditions. Energy minimisation but not equilibration was carried out. The low time step was used because bonds were not constrained. The low γ was used during training only to maximise conformational exploration during the 5 ns simulations. During development gradients were found to be similar when using a γ of 1 ps^-1^. Single precision was used for all floating point values, which gave a significant speedup without a noticeable change in gradient accuracy. Reverse-mode AD has a memory cost proportional to the number of steps in the simulation, which quickly becomes prohibitive. This is alleviated with gradient checkpointing. The simulation is run and the state and random seed are saved every 100 steps; during the reverse pass to get gradients, each block of 100 steps is re-run using the corresponding random seed. Gradient clipping was necessary to prevent gradient explosion. Every 100 steps the norm of the gradients on the coordinates and velocities are calculated, and the gradients are rescaled to have a norm of 0.1 if either norm is greater than 0.1. This is similar to common strategies used to prevent exploding gradients in recurrent neural networks [88]. During development I found that changing the clipping threshold from 0.1 did not have a large effect on the gradients.

Every 5 ps (5000 steps), the Cα residue-residue distances *X* and square distances *X*^2^ are recorded. At the end of the simulation, the mean *µ_s_* = *E*[*X*] and standard deviation 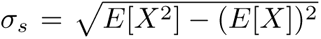 of the Cα residue-residue distances are calculated. For each residue pair, the KL divergence *D_P Q_* to the reference explicit solvent distances (see below) is calculated as

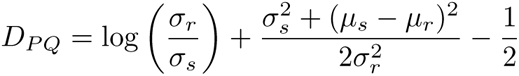

where *µ_r_* and *σ_r_*are the mean and standard deviation of the reference residue pair distances respectively. The KL divergence in the other direction *D_QP_* is also calculated. The loss for each residue pair *L_ij_*is then calculated as

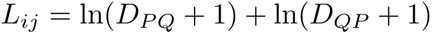

In order to reduce the impact of large losses from residue pairs close in sequence with sharp residueresidue distance distributions, the loss of close residue pairs is downweighted. This meant multiplying *L_ij_* by a weighting factor which is 0 for residue separation *|i − j|* = 0, 1 for *|i − j| ≥* 10 and linearly spaced between. The overall loss is then calculated as the mean of these weighted values over all residue pairs. AD is used to calculate the gradient of this loss with respect to each of the 108 parameters.

At each epoch of training, gradients are combined from simulations of the 8 training proteins to update the force field. For the IDPs, gradients are averaged over the two repeats. For each protein the gradients are then divided by the median of the absolute values of the gradients, meaning that all proteins contribute similarly to the parameter updates each epoch even if the gradient sizes differ. The median was chosen to avoid outliers having too much influence on the value. The gradients for LJ σ parameters and hydrogen parameters were found to be large compared to other parameters and large changes in these parameters can quickly lead to instabilities, so the gradients were weighted by a factor of 0.02. The absolute change for each parameter per protein per epoch was limited to 0.5% and the combined absolute change for each parameter per epoch was limited to 3%. Parameters were updated by gradient descent using a learning rate of 4 *×* 10*^−^*^4^. Training was repeated 3 times and the run with the best performance on the training set was used.

Instead of modifying a partial charge value for each atom type directly, which is complicated by the need to maintain overall charge and by the different partial charges of the same atom type in different residue types, a charge scaling value is learned instead. This starts at one for each atom type. After it is updated during training, partial charges are computed for each atom in a residue type by scaling the starting partial charge by the scaling value and subtracting an offset. This offset is the difference between the starting and scaled overall charge of the residue multiplied by the fraction of the sum of absolute charges present on the atom after scaling. In effect the change in a partial charge of an atom is compensated for by the partial charges of the other atoms in the residue. This allows one charge scaling value to be learned per atom type but the overall charge of each residue type to remain constant during training.

Explicit solvent trajectories used to get reference residue-residue distances for training were generated with Gromacs v2021.4 [86]. For folded proteins the starting structure was the PDB structure and the box size was chosen to give 1 nm padding between the protein and the edge. For disordered proteins the starting structure was the collapsed conformation at the end of a short implicit solvent simulation with a99SB-*disp*+GBNeck2 starting from an extended conformation. The box size was 6 nm for Htt-1-19 and histatin-5 and 7 nm for the two halves of ACTR. These simulations use the a99SB-*disp* force field and its corresponding water model [21], a Verlet leap frog integrator, a 2 fs time step, constrained bonds to hydrogen, a temperature of 300 K, 50 mM NaCl salt, a 1.2 nm cutoff for non-bonded interactions and particle mesh Ewald treatment of long range electrostatics. Energy minimisation, a 100 ps NVT equilibration and a 100 ps NPT equilibration with position restraints to protein heavy atoms preceded a 2 µs production run in the NPT ensemble, with snapshots saved every 50 ps. Mean and standard deviation residue-residue distances were calculated for the last 1 µs of simulation.

Once the force field parameters have been improved via training, simulations can be run with any MD package that supports the GBNeck2 implicit solvent model. Here OpenMM v8.0.0 [85] is used to run the validation simulations as it has high GPU performance and modifying force field parameters is easy. All validation simulations used a Langevin integrator with a γ of 1 ps^-1^, a 4 fs time step, constrained bonds to hydrogen, hydrogen mass repartitioning with a factor of 2, a temperature of 300 K, a 2 nm cutoff for non-bonded interactions and a κ value of 0.7 nm^-1^. Energy minimisation and a 500 ps temperature equilibration with position restraints to heavy atoms preceded production runs, with snapshots saved every 500 ps. Dimers were simulated with no periodic boundaries. Trajectory data was analysed with MDAnalysis [89] and secondary structure was calculated with MDTraj [90]. BioStructures.jl was also used for processing protein structural data [91].

For the simulations of α-synuclein with fasudil, GAFF [92] was used to obtain the force field parameters for fasudil. For each force field, 5 repeats were run starting from different snapshots from α-synuclein monomer simulations. Packmol [93] was used to pack one molecule each of α-synuclein and fasudil in a periodic cubic box with 25 nm sides. A contact is assigned to MD frames where the minimum distance between any fasudil atom and any heavy atom of a residue side chain (CA for glycine) is less than 6 ^°^A [70]. For the Aβ_16-22_ simulations a periodic cubic box with 21.5 nm sides and 6 peptides were used, corresponding to a concentration of 1 mM. The N- and C-termini of the peptides were capped with acetyl (ACE) and *N* -methlyamide (NME) groups respectively to match experimental conditions. Starting conformations for each peptide were taken from snapshots of a short monomer simulation. For each force field and sequence, 3 repeats were run starting from different packings of 6 peptides generated with Packmol. Oligomer size was determined based on groups of contacting peptides, where any pair of atoms being within 4 ^°^A indicates a contact between peptides [73].

## Data and code availability

The trained GB99dms force field in OpenMM XML format, training scripts, simulation scripts and data are available at under a permissive license at https://github.com/greener-group/GB99dms. The Molly.jl software used for training is available at https://github.com/JuliaMolSim/Molly.jl. Simulation trajectories are available at https://zenodo.org/record/8298226.

## Author contributions

Study design, all computational work and manuscript preparation were carried out by JGG.

## Acknowledgements

I thank the Sjors Scheres group and Conny Yu for useful discussions; all contributors to Molly.jl; William Moses and Valentin Churavy for support with Enzyme.jl; Jake Grimmett, Toby Darling and Ivan Clayson for help with high-performance computing; and D. E. Shaw Research for providing data. This work was supported by the Medical Research Council, as part of United Kingdom Research and Innovation (also known as UK Research and Innovation) [MC UP 1201/33]. For the purpose of open access, the MRC Laboratory of Molecular Biology has applied a CC BY public copyright licence to any Author Accepted Manuscript version arising.

## Supplementary Data

**Figure S1.**
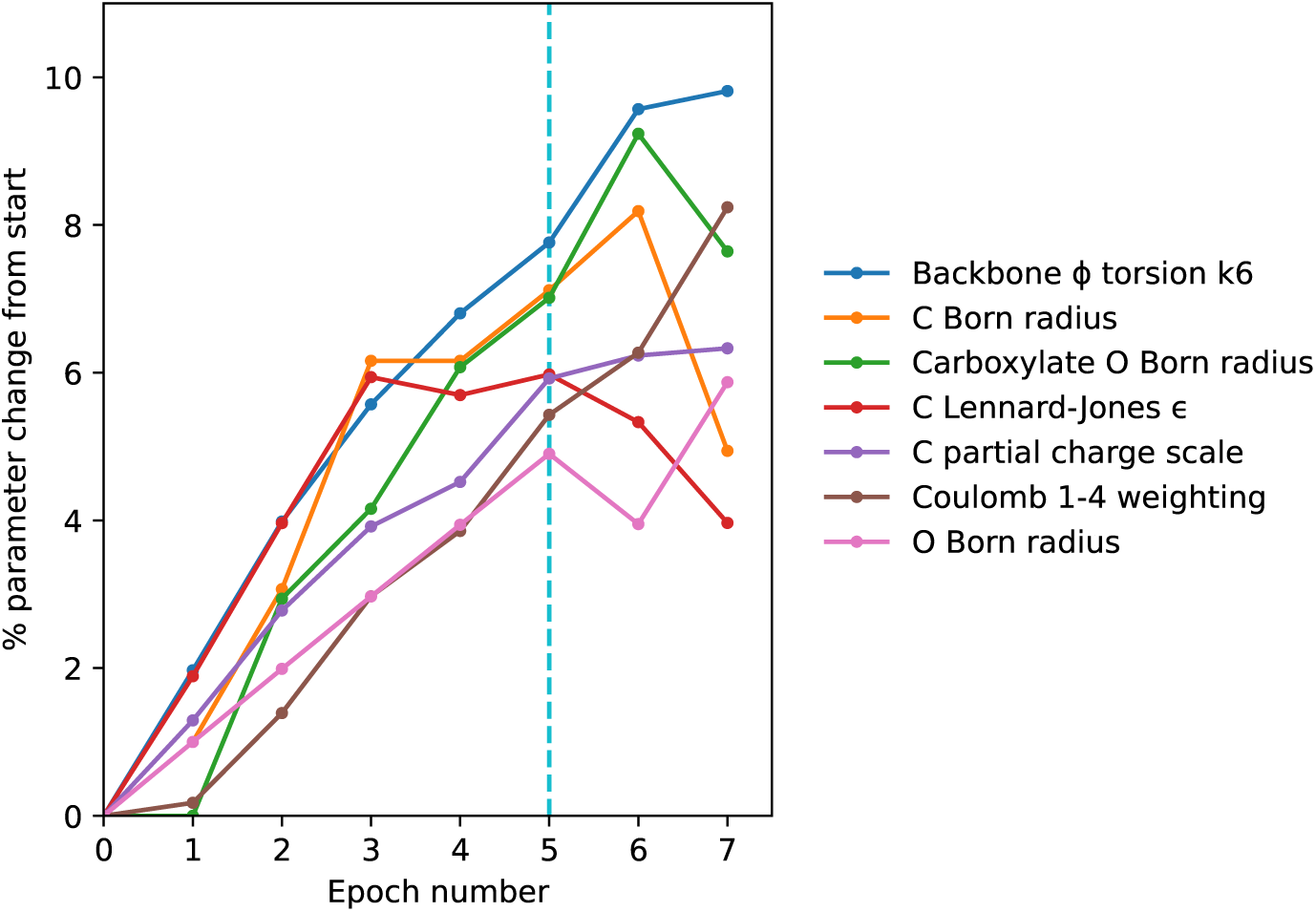
Parameter changes during training. The change in absolute value of 7 parameters that change by at least 4.5% in absolute value from the starting values in a99SB-*disp*+GBNeck2 are shown. Parameters after epoch 5 were used in GB99dms, as indicated by a cyan line.

**Table S1.**
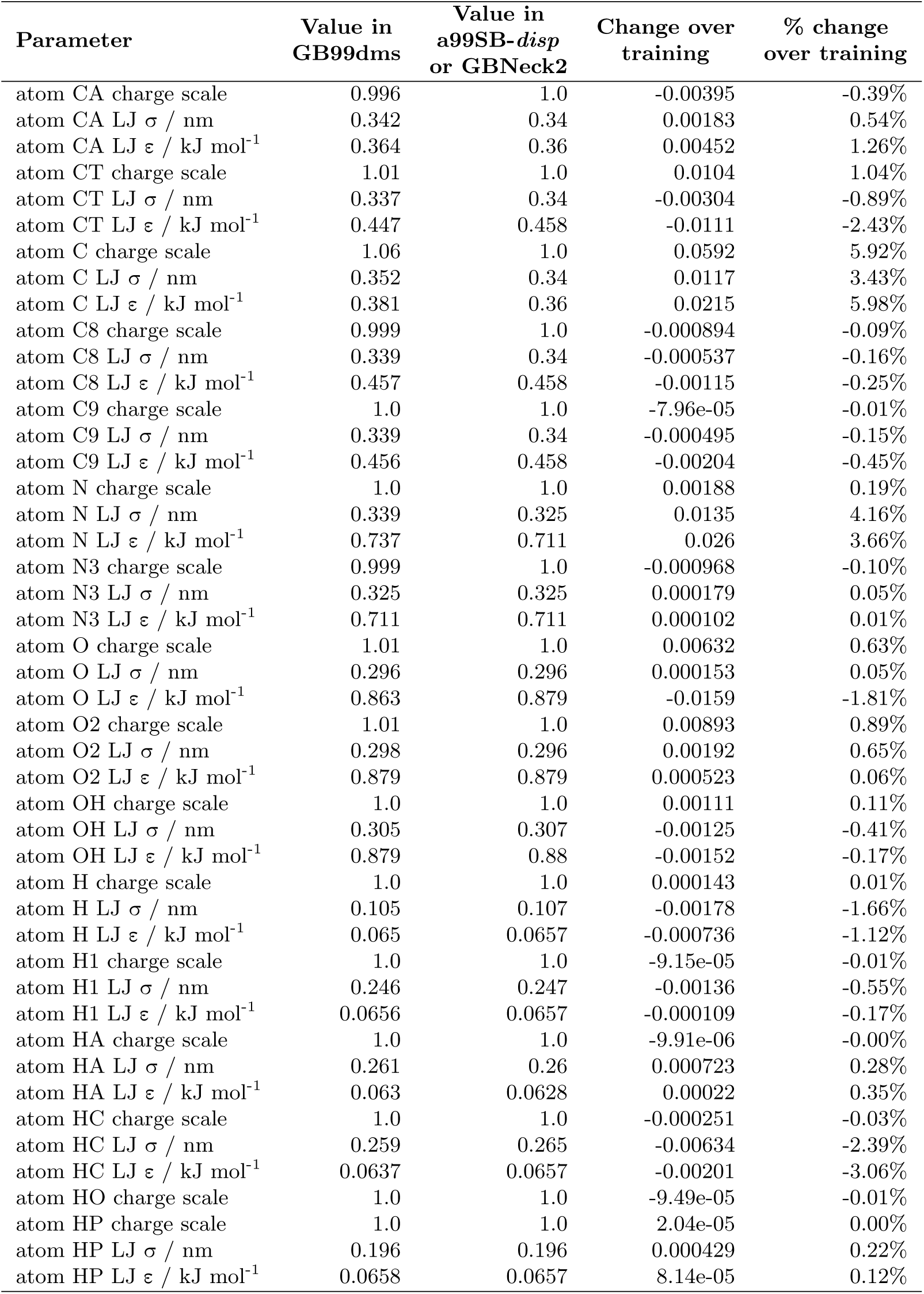
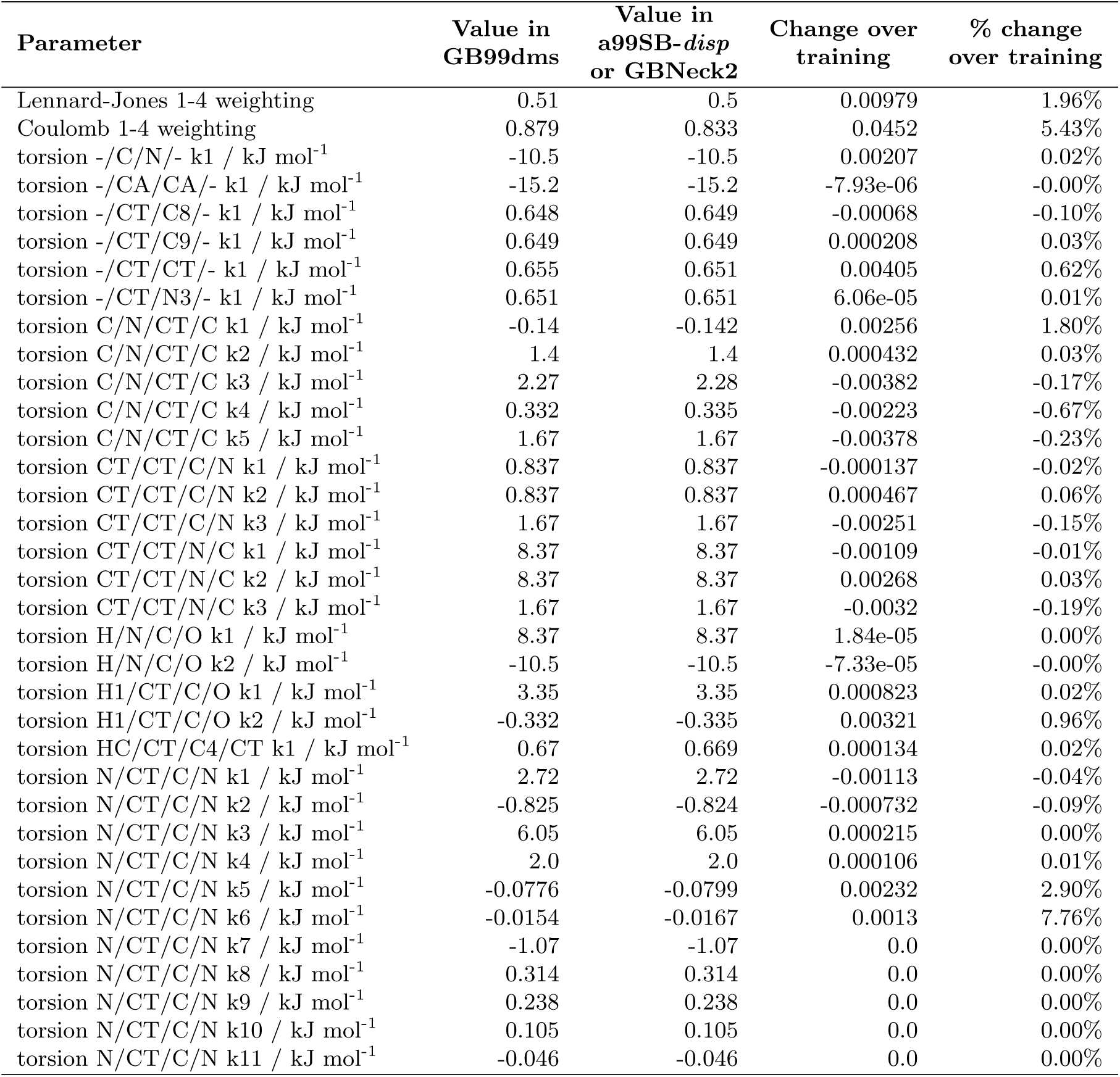
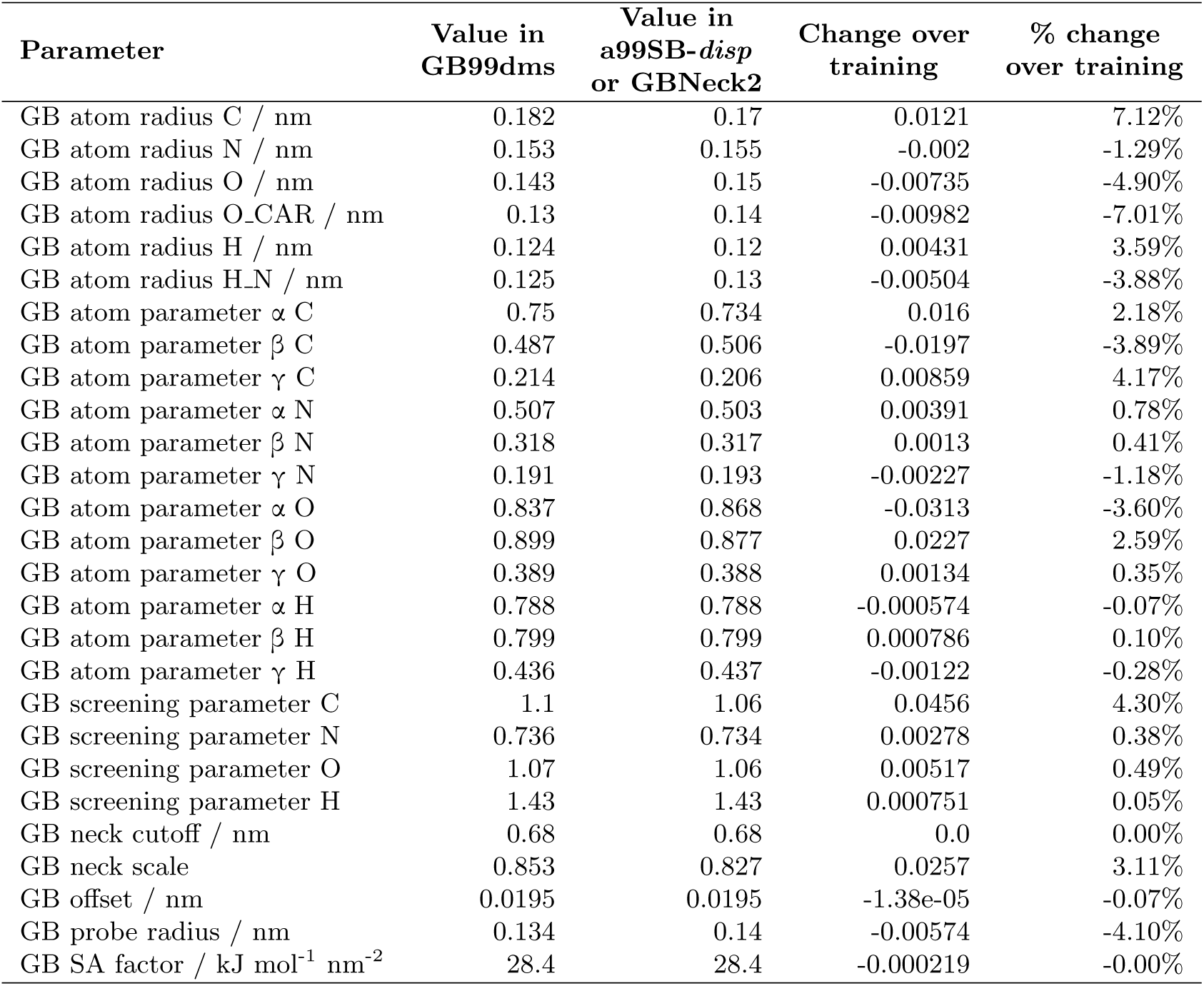
Parameters in GB99dms. The other parameters not changed during training are the same as in a99SB-*disp* or GBNeck2. The atom type HO LJ parameters are zero in a99SB-*disp* so are not modified. This table is available in CSV format in the data repository.

| Parameter | Value in<br>GB99dms | Value in<br>a99SB-*disp*<br>or GBNeck2 | Change over<br>training | % change<br>over training |
| --- | --- | --- | --- | --- |
| atom CA charge scale | 0.996 | 1.0 | -0.00395 | -0.39% |
| atom CA LJ $\sigma$ / nm | 0.342 | 0.34 | 0.00183 | 0.54% |
| atom CA LJ $\epsilon$ / kJ mol<sup>-1</sup> | 0.364 | 0.36 | 0.00452 | 1.26% |
| atom CT charge scale | 1.01 | 1.0 | 0.0104 | 1.04% |
| atom CT LJ $\sigma$ / nm | 0.337 | 0.34 | -0.00304 | -0.89% |
| atom CT LJ $\epsilon$ / kJ mol<sup>-1</sup> | 0.447 | 0.458 | -0.0111 | -2.43% |
| atom C charge scale | 1.06 | 1.0 | 0.0592 | 5.92% |
| atom C LJ $\sigma$ / nm | 0.352 | 0.34 | 0.0117 | 3.43% |
| atom C LJ $\epsilon$ / kJ mol<sup>-1</sup> | 0.381 | 0.36 | 0.0215 | 5.98% |
| atom C8 charge scale | 0.999 | 1.0 | -0.000894 | -0.09% |
| atom C8 LJ $\sigma$ / nm | 0.339 | 0.34 | -0.000537 | -0.16% |
| atom C8 LJ $\epsilon$ / kJ mol<sup>-1</sup> | 0.457 | 0.458 | -0.00115 | -0.25% |
| atom C9 charge scale | 1.0 | 1.0 | -7.96e-05 | -0.01% |
| atom C9 LJ $\sigma$ / nm | 0.339 | 0.34 | -0.000495 | -0.15% |
| atom C9 LJ $\epsilon$ / kJ mol<sup>-1</sup> | 0.456 | 0.458 | -0.00204 | -0.45% |
| atom N charge scale | 1.0 | 1.0 | 0.00188 | 0.19% |
| atom N LJ $\sigma$ / nm | 0.339 | 0.325 | 0.0135 | 4.16% |
| atom N LJ $\epsilon$ / kJ mol<sup>-1</sup> | 0.737 | 0.711 | 0.026 | 3.66% |
| atom N3 charge scale | 0.999 | 1.0 | -0.000968 | -0.10% |
| atom N3 LJ $\sigma$ / nm | 0.325 | 0.325 | 0.000179 | 0.05% |
| atom N3 LJ $\epsilon$ / kJ mol<sup>-1</sup> | 0.711 | 0.711 | 0.000102 | 0.01% |
| atom O charge scale | 1.01 | 1.0 | 0.00632 | 0.63% |
| atom O LJ $\sigma$ / nm | 0.296 | 0.296 | 0.000153 | 0.05% |
| atom O LJ $\epsilon$ / kJ mol<sup>-1</sup> | 0.863 | 0.879 | -0.0159 | -1.81% |
| atom O2 charge scale | 1.01 | 1.0 | 0.00893 | 0.89% |
| atom O2 LJ $\sigma$ / nm | 0.298 | 0.296 | 0.00192 | 0.65% |
| atom O2 LJ $\epsilon$ / kJ mol<sup>-1</sup> | 0.879 | 0.879 | 0.000523 | 0.06% |
| atom OH charge scale | 1.0 | 1.0 | 0.00111 | 0.11% |
| atom OH LJ $\sigma$ / nm | 0.305 | 0.307 | -0.00125 | -0.41% |
| atom OH LJ $\epsilon$ / kJ mol<sup>-1</sup> | 0.879 | 0.88 | -0.00152 | -0.17% |
| atom H charge scale | 1.0 | 1.0 | 0.000143 | 0.01% |
| atom H LJ $\sigma$ / nm | 0.105 | 0.107 | -0.00178 | -1.66% |
| atom H LJ $\epsilon$ / kJ mol<sup>-1</sup> | 0.065 | 0.0657 | -0.000736 | -1.12% |
| atom H1 charge scale | 1.0 | 1.0 | -9.15e-05 | -0.01% |
| atom H1 LJ $\sigma$ / nm | 0.246 | 0.247 | -0.00136 | -0.55% |
| atom H1 LJ $\epsilon$ / kJ mol<sup>-1</sup> | 0.0656 | 0.0657 | -0.000109 | -0.17% |
| atom HA charge scale | 1.0 | 1.0 | -9.91e-06 | -0.00% |
| atom HA LJ $\sigma$ / nm | 0.261 | 0.26 | 0.000723 | 0.28% |
| atom HA LJ $\epsilon$ / kJ mol<sup>-1</sup> | 0.063 | 0.0628 | 0.00022 | 0.35% |
| atom HC charge scale | 1.0 | 1.0 | -0.000251 | -0.03% |
| atom HC LJ $\sigma$ / nm | 0.259 | 0.265 | -0.00634 | -2.39% |
| atom HC LJ $\epsilon$ / kJ mol<sup>-1</sup> | 0.0637 | 0.0657 | -0.00201 | -3.06% |
| atom HO charge scale | 1.0 | 1.0 | -9.49e-05 | -0.01% |
| atom HP charge scale | 1.0 | 1.0 | 2.04e-05 | 0.00% |
| atom HP LJ $\sigma$ / nm | 0.196 | 0.196 | 0.000429 | 0.22% |
| atom HP LJ $\epsilon$ / kJ mol<sup>-1</sup> | 0.0658 | 0.0657 | 8.14e-05 | 0.12% |
| Lennard-Jones 1-4 weighting | 0.51 | 0.5 | 0.00979 | 1.96% |
| Coulomb 1-4 weighting | 0.879 | 0.833 | 0.0452 | 5.43% |
| torsion -/C/N/- k1 / kJ mol<sup>-1</sup> | -10.5 | -10.5 | 0.00207 | 0.02% |
| torsion -/CA/CA/- k1 / kJ mol<sup>-1</sup> | -15.2 | -15.2 | -7.93e-06 | -0.00% |
| torsion -/CT/C8/- k1 / kJ mol<sup>-1</sup> | 0.648 | 0.649 | -0.00068 | -0.10% |
| torsion -/CT/C9/- k1 / kJ mol<sup>-1</sup> | 0.649 | 0.649 | 0.000208 | 0.03% |
| torsion -/CT/CT/- k1 / kJ mol<sup>-1</sup> | 0.655 | 0.651 | 0.00405 | 0.62% |
| torsion -/CT/N3/- k1 / kJ mol<sup>-1</sup> | 0.651 | 0.651 | 6.06e-05 | 0.01% |
| torsion C/N/CT/C k1 / kJ mol<sup>-1</sup> | -0.14 | -0.142 | 0.00256 | 1.80% |
| torsion C/N/CT/C k2 / kJ mol<sup>-1</sup> | 1.4 | 1.4 | 0.000432 | 0.03% |
| torsion C/N/CT/C k3 / kJ mol<sup>-1</sup> | 2.27 | 2.28 | -0.00382 | -0.17% |
| torsion C/N/CT/C k4 / kJ mol<sup>-1</sup> | 0.332 | 0.335 | -0.00223 | -0.67% |
| torsion C/N/CT/C k5 / kJ mol<sup>-1</sup> | 1.67 | 1.67 | -0.00378 | -0.23% |
| torsion CT/CT/C/N k1 / kJ mol<sup>-1</sup> | 0.837 | 0.837 | -0.000137 | -0.02% |
| torsion CT/CT/C/N k2 / kJ mol<sup>-1</sup> | 0.837 | 0.837 | 0.000467 | 0.06% |
| torsion CT/CT/C/N k3 / kJ mol<sup>-1</sup> | 1.67 | 1.67 | -0.00251 | -0.15% |
| torsion CT/CT/N/C k1 / kJ mol<sup>-1</sup> | 8.37 | 8.37 | -0.00109 | -0.01% |
| torsion CT/CT/N/C k2 / kJ mol<sup>-1</sup> | 8.37 | 8.37 | 0.00268 | 0.03% |
| torsion CT/CT/N/C k3 / kJ mol<sup>-1</sup> | 1.67 | 1.67 | -0.0032 | -0.19% |
| torsion H/N/C/O k1 / kJ mol<sup>-1</sup> | 8.37 | 8.37 | 1.84e-05 | 0.00% |
| torsion H/N/C/O k2 / kJ mol<sup>-1</sup> | -10.5 | -10.5 | -7.33e-05 | -0.00% |
| torsion H1/CT/C/O k1 / kJ mol<sup>-1</sup> | 3.35 | 3.35 | 0.000823 | 0.02% |
| torsion H1/CT/C/O k2 / kJ mol<sup>-1</sup> | -0.332 | -0.335 | 0.00321 | 0.96% |
| torsion HC/CT/C4/CT k1 / kJ mol<sup>-1</sup> | 0.67 | 0.669 | 0.000134 | 0.02% |
| torsion N/CT/C/N k1 / kJ mol<sup>-1</sup> | 2.72 | 2.72 | -0.00113 | -0.04% |
| torsion N/CT/C/N k2 / kJ mol<sup>-1</sup> | -0.825 | -0.824 | -0.000732 | -0.09% |
| torsion N/CT/C/N k3 / kJ mol<sup>-1</sup> | 6.05 | 6.05 | 0.000215 | 0.00% |
| torsion N/CT/C/N k4 / kJ mol<sup>-1</sup> | 2.0 | 2.0 | 0.000106 | 0.01% |
| torsion N/CT/C/N k5 / kJ mol<sup>-1</sup> | -0.0776 | -0.0799 | 0.00232 | 2.90% |
| torsion N/CT/C/N k6 / kJ mol<sup>-1</sup> | -0.0154 | -0.0167 | 0.0013 | 7.76% |
| torsion N/CT/C/N k7 / kJ mol<sup>-1</sup> | -1.07 | -1.07 | 0.0 | 0.00% |
| torsion N/CT/C/N k8 / kJ mol<sup>-1</sup> | 0.314 | 0.314 | 0.0 | 0.00% |
| torsion N/CT/C/N k9 / kJ mol<sup>-1</sup> | 0.238 | 0.238 | 0.0 | 0.00% |
| torsion N/CT/C/N k10 / kJ mol<sup>-1</sup> | 0.105 | 0.105 | 0.0 | 0.00% |
| torsion N/CT/C/N k11 / kJ mol<sup>-1</sup> | -0.046 | -0.046 | 0.0 | 0.00% |
| GB atom radius C / nm | 0.182 | 0.17 | 0.0121 | 7.12% |
| GB atom radius N / nm | 0.153 | 0.155 | -0.002 | -1.29% |
| GB atom radius O / nm | 0.143 | 0.15 | -0.00735 | -4.90% |
| GB atom radius O_CAR / nm | 0.13 | 0.14 | -0.00982 | -7.01% |
| GB atom radius H / nm | 0.124 | 0.12 | 0.00431 | 3.59% |
| GB atom radius H_N / nm | 0.125 | 0.13 | -0.00504 | -3.88% |
| GB atom parameter $\alpha$ C | 0.75 | 0.734 | 0.016 | 2.18% |
| GB atom parameter $\beta$ C | 0.487 | 0.506 | -0.0197 | -3.89% |
| GB atom parameter $\gamma$ C | 0.214 | 0.206 | 0.00859 | 4.17% |
| GB atom parameter $\alpha$ N | 0.507 | 0.503 | 0.00391 | 0.78% |
| GB atom parameter $\beta$ N | 0.318 | 0.317 | 0.0013 | 0.41% |
| GB atom parameter $\gamma$ N | 0.191 | 0.193 | -0.00227 | -1.18% |
| GB atom parameter $\alpha$ O | 0.837 | 0.868 | -0.0313 | -3.60% |
| GB atom parameter $\beta$ O | 0.899 | 0.877 | 0.0227 | 2.59% |
| GB atom parameter $\gamma$ O | 0.389 | 0.388 | 0.00134 | 0.35% |
| GB atom parameter $\alpha$ H | 0.788 | 0.788 | -0.000574 | -0.07% |
| GB atom parameter $\beta$ H | 0.799 | 0.799 | 0.000786 | 0.10% |
| GB atom parameter $\gamma$ H | 0.436 | 0.437 | -0.00122 | -0.28% |
| GB screening parameter C | 1.1 | 1.06 | 0.0456 | 4.30% |
| GB screening parameter N | 0.736 | 0.734 | 0.00278 | 0.38% |
| GB screening parameter O | 1.07 | 1.06 | 0.00517 | 0.49% |
| GB screening parameter H | 1.43 | 1.43 | 0.000751 | 0.05% |
| GB neck cutoff / nm | 0.68 | 0.68 | 0.0 | 0.00% |
| GB neck scale | 0.853 | 0.827 | 0.0257 | 3.11% |
| GB offset / nm | 0.0195 | 0.0195 | -1.38e-05 | -0.07% |
| GB probe radius / nm | 0.134 | 0.14 | -0.00574 | -4.10% |
| GB SA factor / kJ mol<sup>-1</sup> nm<sup>-2</sup> | 28.4 | 28.4 | -0.000219 | -0.00% |

**Table S2.** Consistency of gradients. In each case the gradients for the first epoch of training of the run yielding the final model are compared to the gradients for the first epoch of training of the run described. The fraction of corresponding parameter gradients with the same sign and the Pearson correlation coefficient of corresponding parameter gradients are calculated for each protein and the mean over proteins is shown. For the “Noise added to parameters” run, all starting parameter values were multiplied by a number uniformly-distributed in 0.95 to 1.05.

| Run details | Fraction same sign | Correlation |
| --- | --- | --- |
| Repeat 1 | 0.775 | 0.878 |
| Repeat 2 | 0.820 | 0.901 |
| 0.5 fs time step, 2.5 ns simulations | 0.694 | 0.777 |
| Noise added to parameters | 0.767 | 0.859 |

